# Intestinal SEC16B modulates obesity by controlling dietary lipid absorption

**DOI:** 10.1101/2021.12.07.471468

**Authors:** Ruicheng Shi, Wei Lu, Ye Tian, Bo Wang

## Abstract

Genome-wide association studies (GWAS) have identified genetic variants in SEC16 homolog B (*SEC16B*) locus to be associated with obesity and body mass index (BMI) in various populations. *SEC16B* encodes a scaffold protein located at endoplasmic reticulum (ER) exit sites that is implicated to participate in the trafficking of COPII vesicles in mammalian cells. However, the function of SEC16B *in vivo*, especially in lipid metabolism, has not been investigated. Here we demonstrated that intestinal SEC16B is required for dietary lipid absorption in mice. We showed that *Sec16b* intestinal knockout (IKO) mice, especially female mice, were protected from HFD-induced obesity. Loss of SEC16B in intestine dramatically reduced postprandial serum triglyceride output upon intragastric lipid load or during overnight fasting and high-fat diet (HFD) refeeding. Further studies showed that intestinal SEC16B deficiency impaired apoB lipidation and chylomicron secretion. These results revealed that SEC16B plays important roles in dietary lipid absorption, which may shed light on the association between variants in *SEC16B* and obesity in human.

## INTRODUCTION

The prevalence of obesity and its associated metabolic disorders is rising dramatically due to sedentary life style and increased fat consumption (Bluher, 2019; Malik et al., 2013). The intestine plays a central role in lipid absorption, in which lipids are incorporated into triglyceride-rich chylomicrons and transported into circulation (Abumrad and Davidson, 2012; Mansbach and Siddiqi, 2010). It has been shown that elevated chylomicron production contributes to the dyslipidemia in metabolic disorders, such as obesity, diabetes and cardiovascular diseases (Ko et al., 2020). However, the biogenesis and transportation of chylomicron have not been fully understood. In enterocytes, dietary lipids are transferred to apolipoprotein B48 (apoB48) by microsomal triglyceride transfer protein (MTTP) in the endoplasmic reticulum (ER) to form prechylomicrons (Hussain et al., 2003), which are subsequently transported to the Golgi apparatus via prechylomicron transport vesicles (PCTVs) (Siddiqi et al., 2003). The trafficking of PCTVs from the ER to the Golgi involves a cascade of complex processes, including their exit from the ER and delivery to the Golgi (Dash et al., 2015). Their exit from the ER is considered to be the rate-limiting step in chylomicron secretion from enterocyte (Mansbach and Nevin, 1998; Mansbach and Dowell, 2000). Biochemical analysis identified several protein components in PCTVs, such as cluster determinant 36 (CD36), liver-type fatty acid binding protein (L-FABP), and subunits of coat protein II (COPII) machinery (Drover et al., 2005; Nauli et al., 2006; Neeli et al., 2007; Siddiqi et al., 2010; Siddiqi et al., 2003). The function of these proteins in PCTV formation and transportation has not been fully elucidated. Upon fusion to the Golgi, prechylomicrons are further lipidated and secreted into lymphatics as mature chylomicrons.

Genome-wide association studies (GWAS) have identified several single-nucleotide polymorphisms (SNPs) in various genes to be associated with obesity and body mass index (BMI). One of these genes, *SEC16B*, has been linked to obesity in different populations (Albuquerque et al., 2014; Hotta et al., 2009; Lu et al., 2016; Monda et al., 2013; Sahibdeen et al., 2018; Thorleifsson et al., 2009; Vogelezang et al., 2020; Xi et al., 2013; Zeggini et al., 2008). SEC16B is the shorter mammalian orthologue of *S. cerevisiae* SEC16, which was first identified as a scaffold protein that organizes ER exit sites (ERES) by interacting with COPII components (Espenshade et al., 1995; Novick et al., 1980). In mammalian cells, most studies have focused on SEC16A, the longer orthologue that has most similarity to SEC16 in other species. *SEC16A* encodes a 250 kD protein with typical ERES localization (Iinuma et al., 2007; Watson et al., 2006). SEC16A interacts with SEC23 and SEC13 of COPII vesicles and facilitate COPII vesicle budding from the ER, thereby regulating protein secretion (Hughes et al., 2009; Iinuma et al., 2007). In contrast, less is known about the function of SEC16B. *SEC16B* encodes a 117 kD protein that was previously described as a regucalcin gene promoter region-related protein (RGPR-p117) (Yamaguchi, 2009). However, the role of SEC16B as a transcription factor has not been confirmed because there is no evidence that SEC16B directly binds to regucalcin gene promoter region (Tani et al., 2011; Yamaguchi et al., 2003). Compared to SEC16A, SEC16B protein lacks Sec23-interacting C-terminal domain (Bhattacharyya and Glick, 2007). Although both SEC16A and SEC16B play a role in COPII ER exit, several studies indicate that SEC16B likely have specialized, non-redundant functions (Bhattacharyya and Glick, 2007; Budnik et al., 2011). One study demonstrated that SEC16B, but not SEC16A, regulates the transport of peroxisomal membrane biogenesis factor, PEX16, from the ER to peroxisomes in mammalian cells, and thus may participate in the formation of new peroxisomes derived from the ER (Yonekawa et al., 2011). However, the significance of this regulation is unclear. Nevertheless, the pathophysiological functions of SEC16B *in vivo* and whether SEC16B may contribute to the pathogenesis of obesity have not been investigated.

In this study, we examined the role of SEC16B in lipid absorption in the intestine using *Sec16b* intestinal specific knockout (IKO) mice. Our data demonstrate that SEC16B is required for chylomicron production and lipid absorption in the intestine. SEC16B deficiency leads to marked reduction in triglyceride output in serum in the context of bolus lipid challenge. We show that loss of SEC16B impairs chylomicron lipidation and secretion from enterocytes. As a result of lipid malabsorption, *Sec16b* IKO mice are protected from high-fat diet (HFD) induced obesity and glucose intolerance. These data identify SEC16B as a novel regulator of chylomicron production and provide insight into the association between genetic variants in *SEC16B* and obesity in human.

## RESULTS

### *Sec16b* IKO mice do not show noticeable phenotype on chow diet

To examine the function of SEC16B in the intestine, we generated *Sec16b* intestinal specific knockout mice (IKO) from *Sec16b*^tm1a(KOMP)Wtsi^ mice, in which the targeted allele is “conditional-ready”. We crossed heterozygous *Sec16b*^tm1a(KOMP)Wtsi^ mice with mice expressing a *Flpe* transgene to produce a floxed allele. Floxed mice (*Sec16b^F/F^*) were further bred with *Villin-Cre* transgenic mice to delete *Sec16b* exon 13 in the intestine (Fig. 1A). Real-time RT-PCR analysis revealed a ~97% reduction in *Sec16b* mRNA levels in IKO mice compared to floxed *Cre*-negative (F/F) control mice (Fig. 1B). No difference in body weight was observed in either male or female IKO mice compared to their F/F littermates on chow diet (Fig. 1C). Serum triglyceride (TG) and total or free cholesterol levels were comparable between IKO and control in both male and female mice (Fig. S1A-S1C). Interestingly, male IKO mice showed a small but significant decrease in serum non-esterified fatty acid (NEFA) levels compared to control mice. In contrast, female IKO mice displayed a slight increase in serum NEFA levels (Fig. S1D). There was no discernible difference in histology in duodenum and jejunum of IKO intestines compared to control intestines (Fig. 1D).

**Fig. 1.**
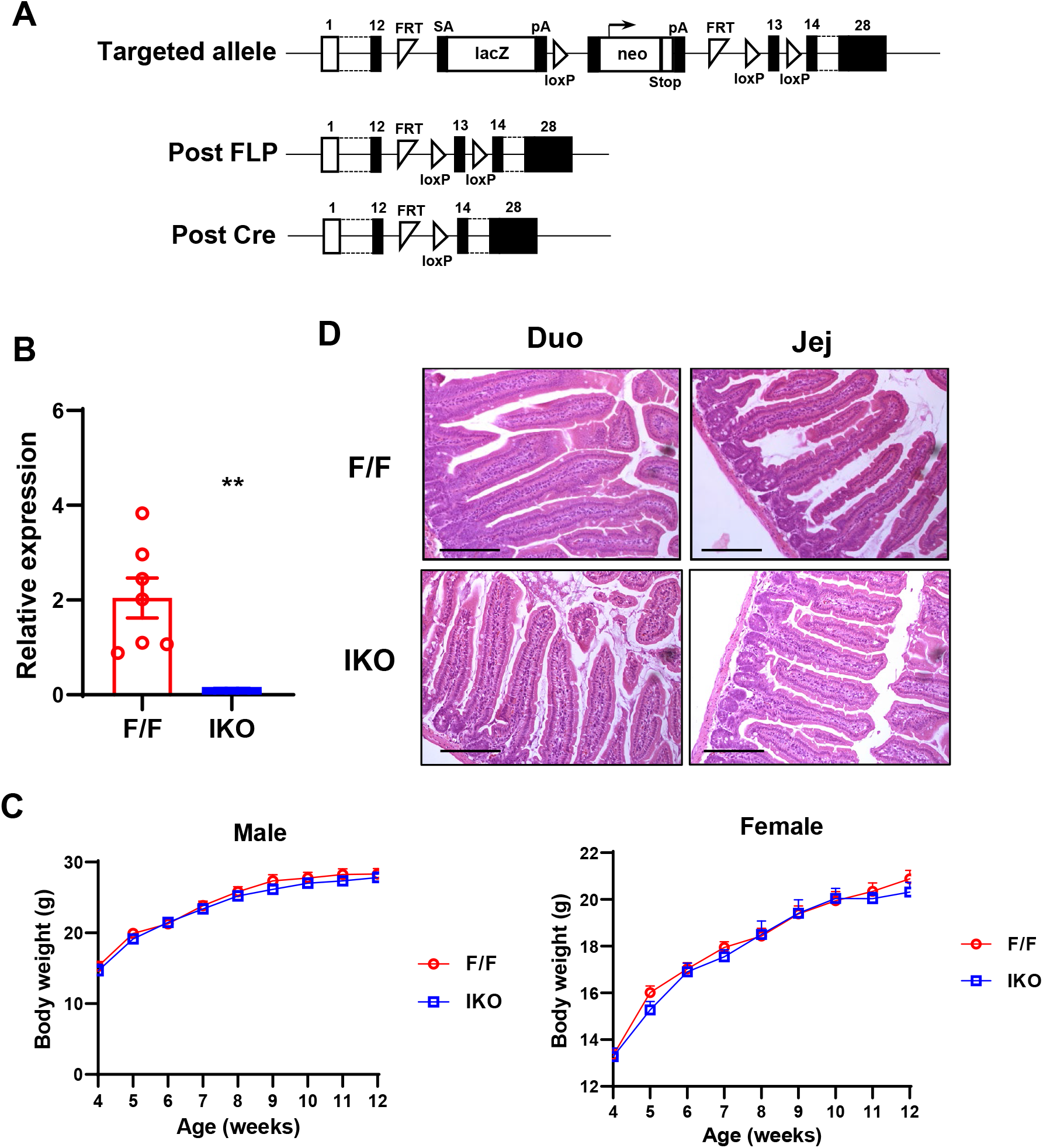
Sec16b IKO mice are normal on chow diet. **A.** Strategy for generating intestine specific Sec16b knockout mice. Sec16b^-/+^ mice carrying the conditional-ready knockout allele were mated with Flpe transgenic mice to generate the Sec16b^F/F^ mice. Sec16b^F/F^ mice were bred with Villin-Cre transgenic mice to generate intestine specific Sec16b knockout mice. **B.** Expression of Sec16b in the intestine of Sec16b^F/F^ (F/F) and Sec16b^F/F^ Villin-Cre (IKO) mice (n= 6-7/group). **C.** Growth curve of Sec16b^F/F^ (F/F) and Sec16b^F/F^ Villin-Cre (IKO) mice on chow diet (n= 6-8/group). **D.** Representative histology of duodenum and jejunum from chow diet fed Sec16b^F/F^ (F/F) and Sec16b^F/F^ Villin-Cre (IKO) mice. Scale bar: 100 μm. Values are means ± SEM. Statistical analysis was performed with Student’s t test. **P <0.01.

### *Sec16b* IKO mice are protected from HFD-induced obesity

To further explore if SEC16B may be involved in the pathogenesis of obesity, we challenged IKO and control mice with HFD starting at 8 weeks of age. Both male and female IKO mice showed significantly less body weight gain compared to control littermates after 12-16 weeks of HFD feeding (Fig. 2A and 2B). Interestingly, female mice showed more pronounced body weight difference between IKO and controls than that in male mice. Magnetic Resonance Imaging (MRI) analysis of body composition showed that the body weight difference after HFD feeding was primarily due to reduced body fat in female mice (Fig. 2C). The body fat mass in male IKO also trended to be less compared to control mice (Fig. 2C). In contrast, lean mass was not changed in both female and male mice (Fig. 2D). Consistent with less obese, both male and female IKO mice showed improved glucose clearance in glucose tolerance test (Fig. 2E and 2F). Serum lipid analysis revealed no difference in TG, NEFA, free cholesterol or cholesterol levels between IKO and control mice after 6 h fasting (Fig. S2A-S2D).

**Fig. 2.**
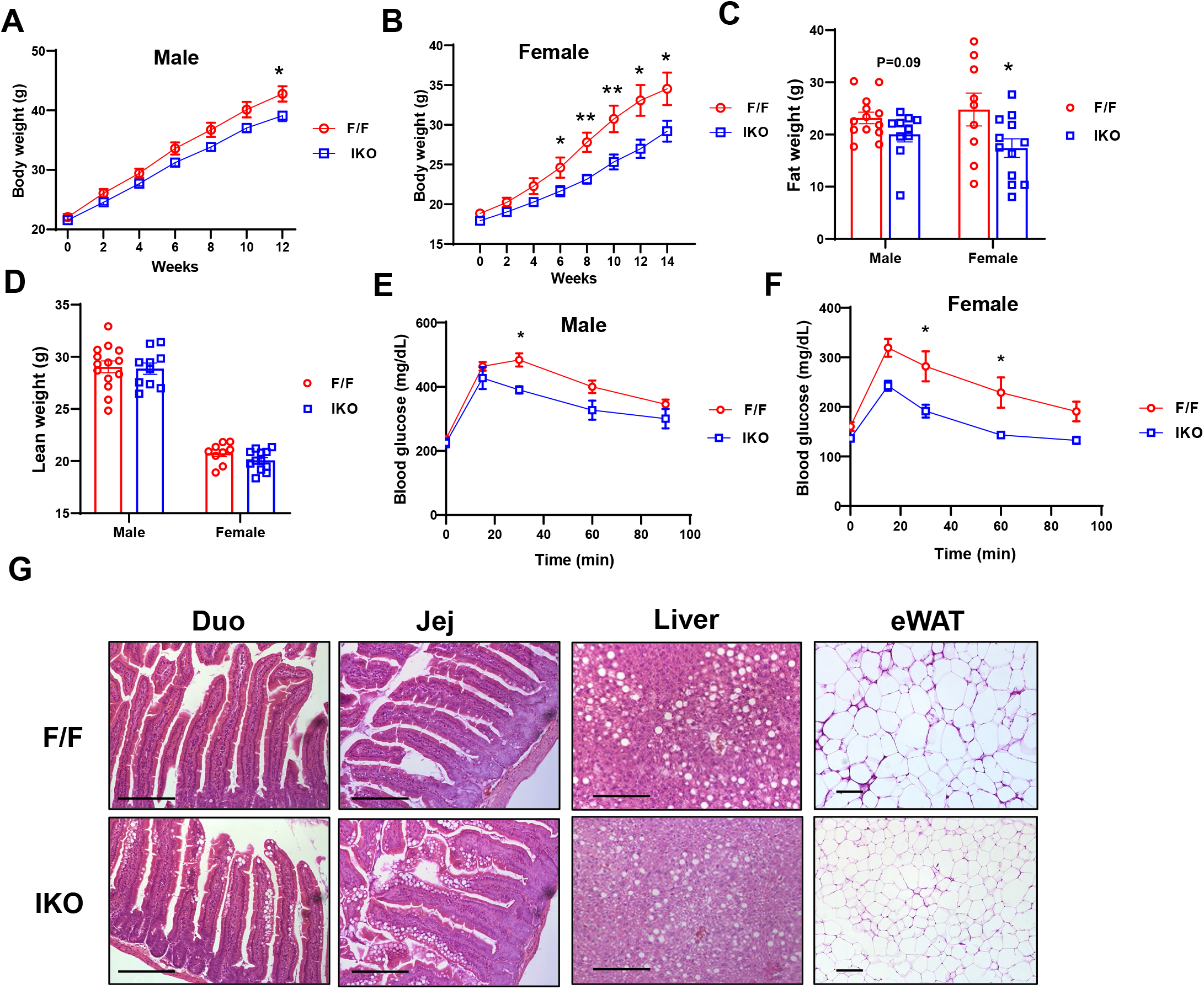
Sec16b IKO mice are protected from HFD-induced obesity. **A** and **B.** Growth curve of male (**A**) and female (**B**) Sec16b^F/F^ (F/F) and Sec16b^F/F^ Villin-Cre (IKO) mice on HFD (n= 9-14/group). **C** and **D.** Fat (**C**) and lean (**D**) mass in HFD-fed Sec16b^F/F^ (F/F) and Sec16b^F/F^ Villin-Cre (IKO) mice (n= 9-14/group). **E** and **F.** GTT analysis of male (**E**) and female (**F**) Sec16b^F/F^ (F/F) and Sec16b^F/F^ Villin-Cre (IKO) mice on HFD (n= 4-8/group). **G.** Representative histology of HFD-fed Sec16b^F/F^ (F/F) and Sec16b^F/F^ Villin-Cre (IKO) mice. Values are means ± SEM. Statistical analysis was performed with Student’s t test (A-D), and two-way ANOVA (E-F). *P < 0.05, **P <0.01.

Next, we sought to understand the cause of reduced body weight gain in IKO mice fed HFD. We first monitored daily food consumption in individually housed mice. *Sec16b* IKO mice consumed a similar amount of food as controls (Fig. S3A), suggesting that this resistance to HFD-induced body weight gain was not due to reduced food consumption in IKO mice. Similarly, there was no difference in fecal TG and NEFA levels between IKO and control mice (Fig. S3B), indicating that loss of SEC16B does not affect lipid uptake from intestinal lumen into enterocytes. Real-time RT-PCR analysis showed that the expression of most of the genes involved in fatty acid (FA) uptake, TG synthesis, lipoprotein production and intracellular transport was not altered in IKO intestines compared to controls (Fig. S4A-S4B). Thus, the expression in lipid metabolic genes could not explain the phenotypes of IKO mice. Next, we measured the energy metabolism in HFD-fed mice by indirect calorimetry using the Comprehensive Lab Animal Monitoring System (CLAMS). As shown in Fig. S5A, there was no difference in oxygen consumption, CO2 production, respiratory exchange ratio (RER) or physical activity in male IKO compared to control mice. In contrast, oxygen consumption and CO2 production trended towards an increase during night cycles in female IKO mice (Fig. S5B), while RER and physical activity were not altered. Thus, increased energy expenditure likely contributes to the more prominent body weight difference in female mice compared to male mice on HFD.

Interestingly, histology analysis of HFD-fed mice revealed more lipid accumulation in IKO intestines compared to controls (Fig. 2G), indicating that chylomicron output and lipid absorption may be impaired in the absence of SEC16B. Hepatic TG, NEFA and total cholesterol levels were comparable between IKO and control mice (Fig. S6A-S6D), while free cholesterol levels were slightly lower in male IKO mice. Consistent with reduced lipid absorption, IKO mice showed much smaller adipocytes and less infiltration of inflammatory cells in epididymal white adipose tissue compared to control mice (Fig. 2G), which may contribute to the improved glucose tolerance in IKO mice. These data demonstrated that loss of SEC16B protects mice from HFD-induced obesity likely through modulating chylomicron metabolism.

### SEC16B deficiency in the intestine impairs lipid absorption

To directly test if SEC16B is required for transport of dietary fat into the circulation, we performed postprandial TG response assays. We gavaged IKO and control mice with a bolus of corn oil and monitored serum TG and NEFA levels at different time points. As expected, serum TG and NEFA levels gradually increased and peaked at 2 h post gavaging in control mice (Fig. 3A and 3B). In contrast, serum TG and NEFA levels remained very low in IKO mice during the entire course. Histology analysis showed marked lipid accumulation in IKO intestines following oil gavage compared to control intestines (Fig. 3C), indicating that loss of SEC16B impairs lipid secretion from enterocytes into circulation. To determine if FA uptake into enterocytes was affected by SEC16B deficiency, we gavaged mice with BODIPY-labeled FA together with corn oil, and examined enterocytes by microscopy. Abundant fluorescently-labeled lipid droplets were observed in IKO intestines, while very little fluorescence was present in controls (Fig. 3D), suggesting that FA uptake was not affected by SEC16B deficiency.

**Fig. 3.**
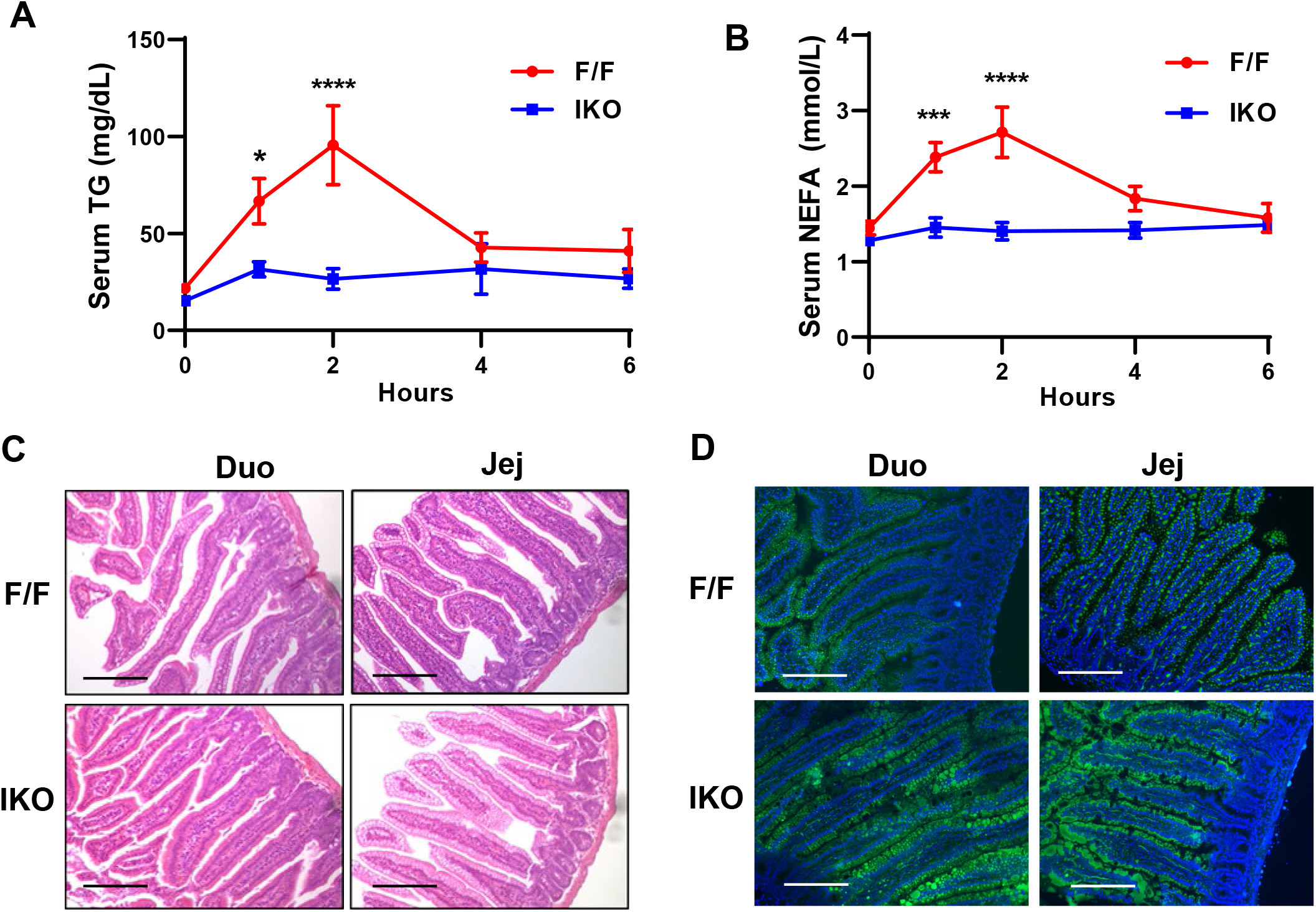
Sec16b deficiency in the intestine impairs lipid absorption when challenged with corn oil bolus. **A** and **B.** Postprandial TG and NEFA response in Sec16b^F/F^ (F/F) and Sec16b^F/F^ Villin-Cre (IKO) mice after oral gavage with corn oil (10 μl/g BW) (n= 9-11/group). **C.** Representative histology of intestine sections from Sec16b^F/F^ (F/F) and Sec16b^F/F^ Villin- Cre (IKO) mice after oral gavage with corn oil for 2 h. Scale bar: 100 μm. **D.** Fluorescence images of small intestines of Sec16b^F/F^ (F/F) and Sec16b^F/F^ Villin-Cre (IKO) mice after oral gavage with olive oil containing BODIPY-labeled fatty acid for 2 h. Scale bar: 100 μm. Values are means ± SEM. Statistical analysis was performed with two-way ANOVA (A, B). *P < 0.05, ***P < 0.001, ****P < 0.0001.

To further determine if intestinal SEC16B regulates lipid absorption, we fasted chow diet fed control and IKO mice overnight and refed them with HFD for 2 h. As shown in Fig. 4A, serum TG levels were ~40% lower in IKO mice compared to controls. Similarly, serum NEFA and free cholesterol levels were significantly decreased in IKO mice, whereas total cholesterol levels were trending towards decrease in IKO mice (Fig. 4B-4D). Consistently, plasma lipoprotein profiling analysis revealed a marked decrease in TG levels in chylomicron/VLDL fraction in IKO mice upon fasting/HFD refeeding (Fig. 4E), while cholesterol levels were decreased in all lipoprotein fractions in IKO serum (Fig. 4F). Similar to oil gavage, HFD refeeding resulted in massive lipid accumulation in both duodenum and jejunum of IKO mice as revealed by histology analysis and Oil red O staining (Fig. 4G and 4H). Taken together, these data demonstrated that loss of SEC16B in the intestine impairs lipid absorption likely through blunting lipid secretion from enterocytes into the circulation without affecting FA uptake.

**Fig. 4.**
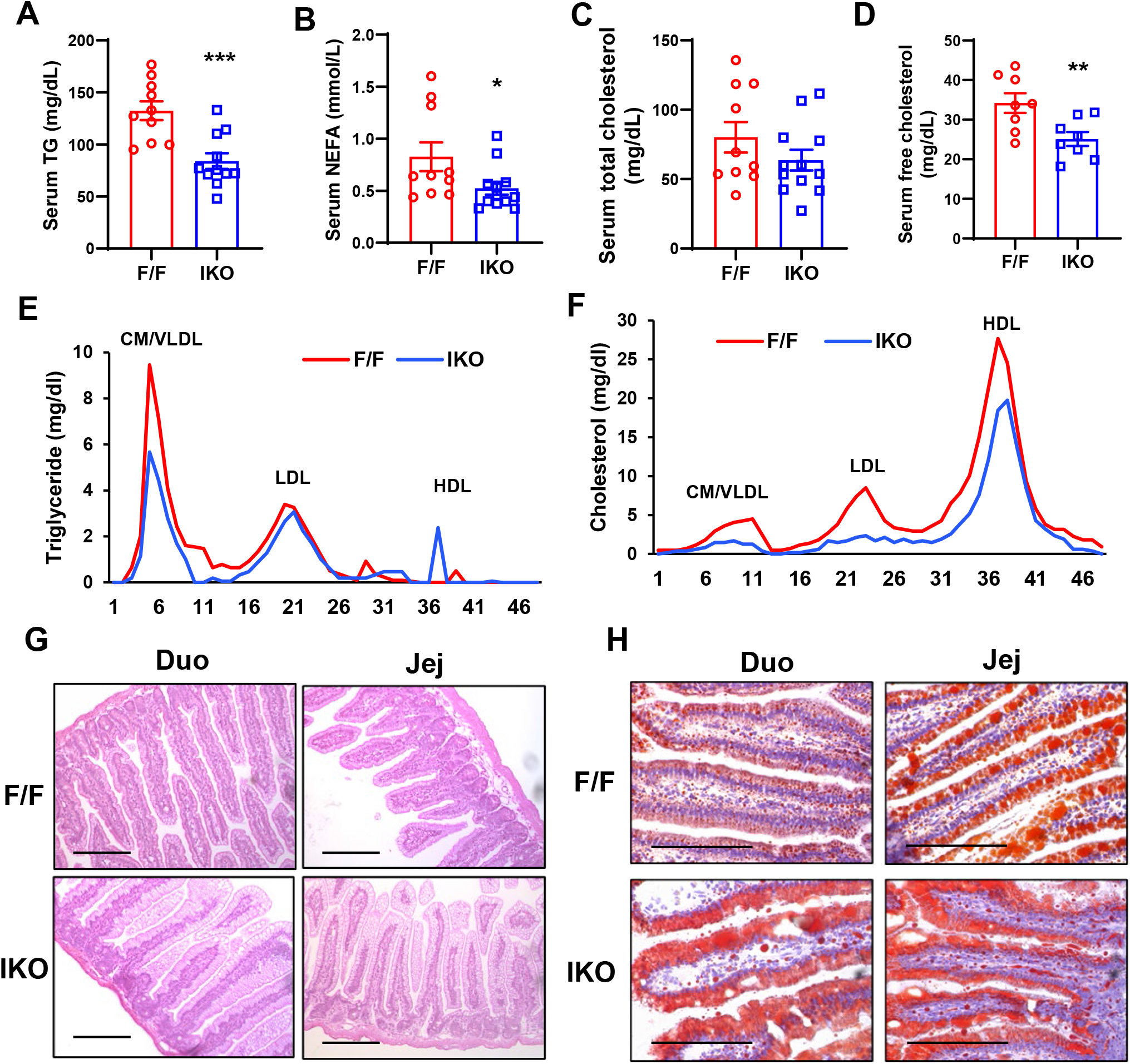
Loss of Sec16b in the intestine impairs lipid absorption during fasting and HFD refeeding. **A-D.** Plasma lipid levels in chow diet fed Sec16b^F/F^ (F/F) and Sec16b^F/F^ Villin-Cre (IKO) mice after fasting overnight and refeeding HFD for 2 h (n= 8-12/group). **E-F.** Plasma from *Lpcat3^fl/fl^* (F/F) and *Lpcat3^fl/fl^ Albumin-Cre* (L-Lpcat3-KO) mice fasted overnight was pooled (*n* = 5). Lipoprotein profiles were analyzed by fast protein liquid chromatography (FPLC). **G.** Representative histology of intestine sections from Sec16b^F/F^ (F/F) and Sec16b^F/F^ Villin-Cre (IKO) mice as in A-D. Scale bar: 100 μm. **H.** Representative oil-red-O staining of intestine sections from Sec16b^F/F^ (F/F) and Sec16b^F/F^ Villin-Cre (IKO) mice as in A-D. Scale bar: 100 μm. Values are means ± SEM. Statistical analysis was performed with Student’s t test (A-D). *P < 0.05, **P < 0.01, ***P < 0.001.

### Loss of SEC16B impairs chylomicron lipidation and secretion

To understand how loss of SEC16B affects lipid absorption, we first examined the expression of genes involved in lipid metabolism in the intestines of overnight fasting and HFD refeeding mice. The mRNA levels of acyl-CoA synthase long chain family member 5 (*Acsl5*) and monoacylglycerol O-acyltransferase 2 (*Mogat2*), two genes in the TG synthesis pathway in enterocytes, were significantly decreased in male IKO jejunum compared to controls after overnight fasting (Fig. S7A). There was also a trend of decrease in the expression of *Mttp* (P=0.06) in male IKO intestines. However, the changes in their expression were abolished upon HFD refeeding (Fig. S7B). Furthermore, there was no significant change in the expression of these genes in female IKO mice compared to controls in either fasting or refeeding conditions (Fig. S7C-S7D). These data indicate that reduced lipid absorption in IKO mice is unlikely caused by altered expression of lipid metabolic genes.

Dietary lipids are absorbed into circulation in the form of chylomicron. We next investigated if intestinal SEC16B deficiency affects chylomicron assembly and secretion upon HFD refeeding. Negative staining of plasma chylomicron fractions by electron microscopy revealed markedly smaller chylomicron particles in *Sec16b* IKO mice compared to controls (Fig. 5A and 5B), suggesting poor apoB lipidation. ApoB levels in the chylomicron fraction were decreased in *Sec16b* IKO mice, while total serum apoB levels were comparable (Fig. 5C), indicating that chylomicron secretion is impaired in the setting of SEC16B deficiency. Interestingly, we did not observe accumulation of apoB48 in the intestine of IKO mice (Fig. 5D), likely because its degradation was enhanced due to poor lipidation. Despite reduced apoB lipidation, MTTP protein levels were comparable in IKO and control mice after 2 h of HFD refeeding (Fig. 5D). To further investigate if SEC16B affects PCTV trafficking within enterocytes, we isolated ER and Golgi fractions from enterocytes of HFD refed mice, and analyzed the distribution of apoB in ER and Golgi. As shown in Fig. 5E, apoB level was slightly higher in ER fraction, but much lower in Golgi fraction of IKO enterocytes compared to controls, indicating that loss of SEC16B likely blunts the transport of PCTV from ER to Golgi.

**Fig. 5.**
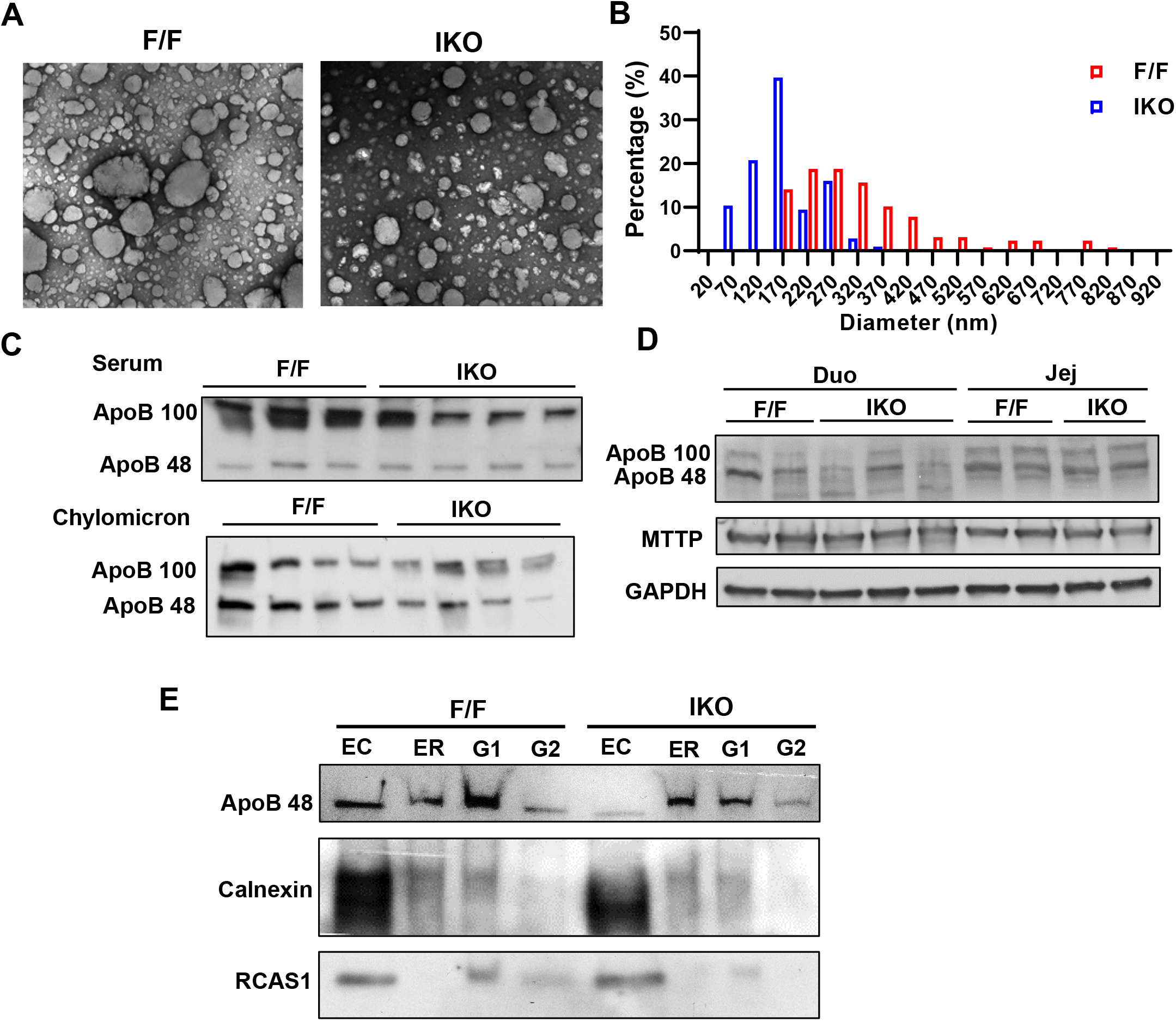
Sec16b deficiency in the intestine impairs chylomicron lipidation and secretion. **A.** Plasma chylomicron particle size in Sec16b^F/F^ (F/F) and Sec16b^F/F^ Villin-Cre (IKO) mice. Chow diet fed mice were fasted overnight and refed 60% HFD for 2 h. Chylomicrons were isolated and pooled from 5 mice/group. Chylomicrons were stained with 2.0% uranyl acetate and visualized by electron microscopy. **B.** Quantification of chylomicron particle size in (A). **C.** ApoB western blot in whole serum and chylomicron fraction of Sec16b^F/F^ (F/F) and Sec16b^F/F^ Villin-Cre (IKO) mice as in (A). **D.** ApoB western blot in duodenum and jejunum of Sec16b^F/F^ (F/F) and Sec16b^F/F^ Villin-Cre (IKO) mice as in (A). **E.** ApoB western blot in endoplasmic reticulum (ER) and Golgi isolated from the enterocytes of Sec16b^F/F^(F/F) and Sec16b^F/F^ Villin-Cre (IKO) mice as in (A). EC: enterocyte whole cell lysate, G1: Golgi fraction #1, G2: Golgi fraction #2.

To obtain further insight into the mechanism of the chylomicron production defect in *Sec16b* IKO mice, we examined intestine samples by electron microscopy (EM). Consistent with more lipid accumulation in HFD refed IKO intestines, EM analysis showed much bigger cytosolic lipid droplets (CLDs) in IKO enterocytes (Fig. 6A). In contrast, IKO enterocytes contained markedly smaller lipoprotein particles in secretory vesicles compared to control enterocytes (Fig. 6B). Consistently, lipoprotein particles were almost undetectable in the intercellular space of IKO intestines (Fig. 6C). These data suggest that SEC16B regulates enterocyte TG distribution and chylomicron transport in enterocytes.

**Fig. 6.**
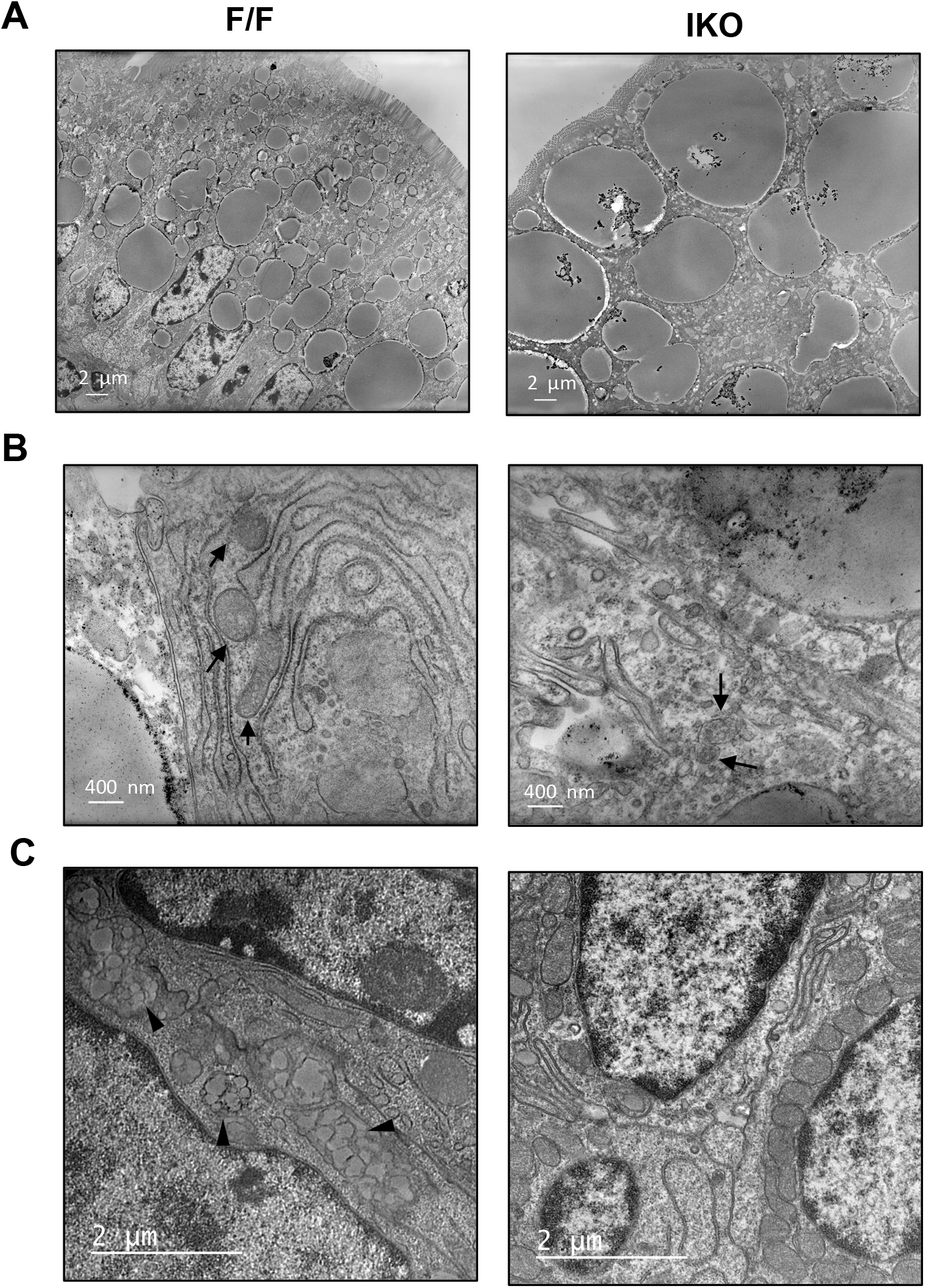
Reduced chylomicron secretion in Sec16b deficient enterocytes. **A-C.** Electron microscopy analysis of intestine sections from Sec16b^F/F^ (F/F) and Sec16b^F/F^ Villin-Cre (IKO). Chow diet fed mice were fasted overnight and refed 60% HFD for 2 h. Arrows indicate secretory vesicles containing lipoprotein particles. Arrowheads indicate chylomicron in the chylomicron in the intercellular space.

## DISCUSSION

The small intestine plays an important role in modulating lipid homeostasis by controlling lipid absorption. Excess lipid absorption is one of the major risk factors of obesity and metabolic disorders (Bluher, 2019; Klop et al., 2013). The absorption of dietary lipids involves several steps, including the uptake of FA and monoacylglycerol from intestinal lumen, resynthesis of TG, chylomicron assembly, lipidation and secretion(Abumrad and Davidson, 2012; Dash et al., 2015; Mansbach and Siddiqi, 2010). Although the molecular mechanisms modulating these steps have been extensively investigated, the regulation of chylomicron transport and secretion in enterocytes remains poorly understood. Here we have shown that loss of SEC16B in intestine impairs lipid absorption likely through blocking chylomicron transport from ER to Golgi and impeding the lipidation of prechylomicrons. These findings implicate that SEC16B is a critical regulator of chylomicron maturation and secretion.

Previous studies have shown that Sec16 is localized at transitional ER (tER), where they function as scaffold proteins to direct the organization of ERES in yeast and *Drosophila* (Connerly et al., 2005; Espenshade et al., 1995; Ivan et al., 2008). Sec16 interacts with Sec23, Sec24 and Sec31 in COPII machinery and control its exit from ER to Golgi in the secretory pathway (Gimeno et al., 1996; Shaywitz et al., 1997). It is well documented that COPII proteins are involved in the transport of proteins and lipoprotein particles, including chylomicrons (Jones et al., 2003; Mansbach and Siddiqi, 2010; Zanetti et al., 2011). Therefore, Sec16 has been proposed to regulate protein and lipid transport. Several studies demonstrated that mutations in Sec16 gene result in the accumulation of secretory proteins and lack of secretory vesicles in the cell (Bhattacharyya and Glick, 2007; Espenshade et al., 1995; Kaiser and Schekman, 1990; Watson et al., 2006), corroborating that Sec16 participates in protein secretion. However, whether Sec16 modulates lipid secretion has not been investigated. Mammalian cells contain two Sec16 homologs, SEC16A and SEC16B, which both localize at ERES (Bhattacharyya and Glick, 2007). SEC16A is the primary homolog of yeast Sec16 as they have similar molecular mass (Iinuma et al., 2007). *In vitro* studies have shown that SEC16A plays a major role in the organization of ERES and protein export from the ER (Bhattacharyya and Glick, 2007; Iinuma et al., 2007). In contrast, SEC16B plays a minor role in protein secretion as depletion of SEC16B has a lesser effect compared to loss of SEC16A (Tani et al., 2011). Moreover, overexpression of SEC16B cannot compensate the defect in protein secretion caused by SEC16A knockdown (Budnik et al., 2011). Furthermore, we showed that loss of SEC16B in intestine does not result in discernible phenotype on chow diet despite the fact that small intestine is an important endocrine organ that secrets gut hormones, indicating that SEC16B is unlikely to play critical roles in protein secretion.

Interestingly, SEC16A and SEC16B show different tissue distribution in mammals. SEC16A is ubiquitously expression in most tissues, while SEC16B is highly expressed in the liver and intestine (data not shown), suggesting that they may have distinct functions in these organs. Considering that the liver and small intestine are the major organs for lipoprotein production in the body, we hypothesized that SEC16B may be involved in lipoprotein metabolism. Our data showed that SEC16B deficiency in intestine impairs lipid absorption, supporting a critical role in chylomicron metabolism. In addition, we did not observe compensatory effect in defective lipid absorption by endogenous SEC16A in IKO mice, suggesting that the function in regulating lipid metabolism is likely unique to SEC16B. But how SEC16B acquires this specialized function during evolution is still unclear. Compared to SEC16A, SEC16B lacks a C-terminal conserved domain that interacts with other proteins (Bhattacharyya and Glick, 2007). Thus, it is possible that SEC16B interacts with a different set of proteins in metabolic tissues *in vivo* to fulfill its function in lipid metabolism.

Our mechanistic studies revealed that loss of SEC16B blunts lipid absorption through blocking its transport from ER to Golgi and inhibiting chylomicron lipidation. Given that SEC16B localizes to ERES and interacts with COP II components, it is rational to speculate that it may regulate COP II exit from ER, and thereby modulating chylomicron trafficking. But the observation that IKO mice have reduced chylomicron lipidation is surprising since there was no difference in MTTP protein levels in control and IKO intestines. Interestingly, IKO enterocytes contained much bigger cytosolic lipid droplets compared to controls. Considering that the biogenesis of both lipid droplets and chylomicrons are initiated in the ER, we speculate that lipids may be channeled to lipid droplets instead of being incorporated into chylomicron particles in IKO mice. Further studies will be needed to determine if SEC16B is involved in the partitioning of lipids to lipid droplets and chylomicrons.

Numerous GWAS studies have demonstrated that several SNPs in *SEC16B* gene locus are associated with obesity and BMI. For examples, the minor “C” variant of rs10913469 in the intron of *SEC16B* is more common in adults and children with obesity and higher BMI in Asia, Mexico, and several European countries (Hotta et al., 2009; Jimenez-Osorio et al., 2019; Thorleifsson et al., 2009; Zhao et al., 2014). Similarly, the intergenic “G” variant of rs543874 is associated with higher obesity rate and body fat percentage in Africa American, East Asian and European descent (Lu et al., 2016; Monda et al., 2013; Vogelezang et al., 2020). However, it is unclear how these variants may affect body weight and fat mass in the body. Our studies suggest that SEC16B likely contributes to the pathogenesis of obesity through regulating lipid absorption. It would be interesting to examine if these variants affect the expression of SEC16B and whether overexpression of SEC16B increases lipid absorption and adiposity. Interestingly, genetic variants in *SEC16B* have been shown to be associated with obesity more significantly in women (Shi et al., 2010; Xi et al., 2013), which is consistent with our observation that the difference in body weight gain on HFD was more dramatic in female IKO mice than males. The mechanisms underlying gender difference in the roles of SEC16B in obesity merits further investigation.

Nevertheless, these results revealed a novel regulator of the biogenesis and transport of chylomicron, and thereby controlling lipid absorption and obesity. Future studies will explore whether targeting SEC16B could be used as a strategy to modulate diet-induced obesity.

## MATERIAL AND METHODS

### Generation of Sec16b IKO mice

We generated *Sec16b* intestinal specific knockout mice (IKO) from *Sec16b*^tm1a(KOMP)Wtsi^ (MMRRC_049583_UCD) mice, in which the targeted allele is “conditional-ready”. We first crossed heterozygous *Sec16b*^tm1a(KOMP)Wtsi^ mice with mice expressing a *Flpe* recombinase deleter transgene (B6.Cg-Tg(ACTFLPe)9205Dym/J) to remove the gene-trapping cassette in intron 12 of *Sec16b*, producing a conditional knockout allele (*Sec16b^F/+^*) containing loxP sites in intron 12 and intron 13. *Sec16b^F/F^* mice were crossed with *villin-Cre* transgenic mice (B6.Cg-Tg(Vil1-cre)997Gum/J) to generate *Sec16b^F/F^, Villin-Cre* + (IKO) mice. *Sec16b^F/F^, Villin-Cre* – mice were used as controls.

### Mouse studies

All animal procedures were conducted in compliance with protocols approved by the Institutional Animal Care and Use Committee (IACUC) at University of Illinois at Urbana-Champaign (UIUC). All mice were housed under pathogen-free conditions in a temperature-controlled room with a 12 h light/dark cycle. All mice were subjected to either chow diet, HFD (60% calories from fat, Research Diets #D12492). All mice were fasted for 6 h prior experiments unless stated otherwise. Small intestine samples were flushed with cold PBS and collected at different regions (duodenum, jejunum and ileum) corresponding to a length ratio of 1:2:1. All samples were snap frozen in liquid nitrogen and stored in −80°C, or fixed in 10% formalin, or frozen in OCT for cryosectioning. Blood was collected by retro-orbital bleeding, and the plasma was separated by centrifugation. Plasma lipids were measured with the Wako Free Cholesterol E kit, the Wako Cholesterol E kit, the Wako HR series NEFA-HR (2) kit (FUJIFILM, Richmond, VA), and the Infinity Triglyceride Reagent kit (Thermo Fisher, Waltham, MA). Tissue and fecal lipids were extracted with Folch lipid extraction (Folch et al., 1957) and measured with the same enzymatic kits. For glucose tolerance tests (GTT), mice were fasted for 6 h and i.p. injected with D-Glucose (1 g/kg body weight), blood glucose levels were measured at 0, 15, 30, 60 and 90 mins. Tissue histology was performed in the UIUC Comparative Biosciences Histology Laboratory.

### Imaging Studies

Mice were fasted for 4 h and gavaged with BODIPY™ 500/510 C1, C12 (2 μg/g body weight) in corn oil (10 μl/g body weight). After 2 h, small intestines (duodenum and jejunum) were harvested and embedded in OCT. Samples were sectioned into 10 μm sections, immediately mounted in ProLong^TM^ Diamond Antifade Mountant with DAPI and examined under fluorescence microscope. For fasting and HFD refeeding experiment, mice were fasted overnight and refed with HFD for 2 h. Duodenum and jejunum were harvested, embedded, and sectioned as described above. Slides were then fixed in 4% paraformaldehyde for 5 min, stained in Oil-Red-O for 10 min and counterstained with hematoxylin. For determination of chylomicron size by transmission electron microscopy (TEM), 100 μl pooled plasma from 4 *Sec16b* IKO and *Sec16b^F/F^* control mice were overlaid with 700 μl saline and centrifuged at 70, 000 rpm for 5 h in a TLA 120.2 rotor and the top layer was collected. For electron microscopy analysis, 5 μl of the chylomicron fraction was applied to carbon-coated copper grids and stained with 2.0% uranyl acetate for 15 min. Grids were visualized with a JEOL 100CX transmission electron microscope. Particle diameter was measured using ImageJ.

### Transmission electron microscopy analysis for intestine

Control and IKO mice were fasted overnight and refed HFD for 2 h before sacrificing. Upon refeeding, mouse tissues (duodenum and jejunum) were collected and fixed immediately in Sorenson’s Phosphate Buffer mixture containing 2% paraformaldehyde and 2.5% glutaraldehyde. The samples were then rinsed in phosphate buffer, incubated in Osmium tetroxide with potassium ferrocyanide, and rinsed with water. Samples were en-bloc stained with filtered uranyl acetate overnight and dehydrated in ethanol series. Samples were then subsequently infiltrated with 1:1 and 1:4 acetonitrile: epoxy mixture and finally infiltrated with Lx112 epoxy mixture (Ladd, Inc), and hardened at 80 °C for 2 days. After embedding, samples were trimmed, and cut into 90-100 nm sections with a diamond knife. Sections were stained with Uranyl Acetate and Lead Citrate and were visualized on a Hitachi H600 Electron Microscope at 75 KV. Images were taken with plate film and scanned at 3200 dpi for digital images.

### Postprandial lipid absorption assay

For lipid absorption assay, mice were fasted for 4 h and gavaged with corn oil (10 μg/g body weight). Plasma was collected through tail vein at 0, 1, 2, 4 and 6 h and plasma lipids were measured with Wako HR series NEFA-HR (2) kit and the Infinity Triglyceride Reagent kit. Plasma fast protein liquid chromatography (FPLC) lipoprotein profiles were performed at Lipid Core of Vanderbilt University School of Medicine.

### Endoplasmic reticulum and Golgi fractionation from enterocyte

The ER and Golgi fractions were isolated from enterocytes as described (Siddiqi et al., 2003). In brief, enterocytes were harvested from overnight fasting and HFD refed control and IKO mice following previous publication (Wang et al., 2016). 2×10^7^ cells were lysed in buffer A (0.25 M Sucrose, 5 mM EDTA, 10 mM HEPES, protease inhibitor cocktail) and homogenized using a motor driven homogenizer (Caframo, Ontario, Canada). The homogenate was spun at 8,500 g for 10 min at 4°C, and the resulting supernatant was further spun at 100,000g for 3 h at 4°C. The pellets were collected, density adjusted with 1.22 ml 1.22M sucrose solution, the resuspended mixture was then overlaid with 1 ml 1.15 M, 1 ml 0.86 M and 0.7 ml 0.25 M sucrose solution. This discontinuous gradient was spun at 82,000g for 3 h at 4°C. Golgi fraction was enriched at 0.25/0.86 and 0.86/1.15 interface, while ER/microsomal membrane remained in the 1.22 M sucrose layer and/or appeared as a pellet. All fractions were collected with the same volume.

### Quantitative PCR

In brief, tissue was homogenized with TissueLyser II (Qiagen, Hilden, Germany), and total RNA was extracted with TRIzol (Invitrogen, Waltham, MA). cDNA was synthesized, and gene expression was quantified by CFX384 Touch Real-Time PCR Detection System (Bio-Rad, Hercules, CA) with SYBR Green (Thermo Fisher, Waltham, MA). Gene expression levels were normalized to 36B4.

### Western Blot

Intestine tissue was homogenized by TissueLyser II in RIPA buffer (50 mM Tris–HCl, pH 7.4, 150 mM NaCl, 1% NP-40, 0.5% sodium deoxycholate, 0.1% SDS) supplemented with protease and phosphatase inhibitors and PMSF. Lysates were sonicated and supernatant was collected by centrifugation. Protein lysates, sera, or chylomicron fractions were mixed with 4x laemmli buffer and loaded onto 4%–15% TGX Gels (Bio-Rad, Hercules, CA), transferred to hybond PVDF membrane (GE Healthcare, Chicago, IL), and incubated with the following antibodies: anti-APOB (ab20737, Abcam), anti-MTTP (sc-135994, Santa Cruz Biotechnology), GAPDH (MAB374, EMD Millipore). After incubation with secondary antibodies, the protein bands were visualized with enhanced chemiluminescence (ECL) (Thermo Fisher, Waltham, MA).

### Indirect Calorimetry and Body Composition Measurements

Metabolic rates were measured by indirect calorimetry in open-circuit Oxymax chambers in the Comprehensive Lab Animal Monitoring System (CLAMS) (Columbus Instruments). Mice fed with HFD for 10-12 weeks were placed individually in metabolic chambers of the CLAMS (Columbus Instruments, Columbus, OH) for one week. The chamber was maintained at 23°C with 12 h light/dark cycles. Food and water were provided as needed. Oxygen consumption rate (VO_2_), carbon dioxide production (VCO_2_) rates, respiratory exchange ratio (RER) and physical activities (x-tot) were measured every 12 minutes over a period of 2 days. The first readings were taken after a 36 h acclimation period and data were presented as a 3 day average line chart. Body compositions (whole-body fat and lean composition) were measured on Echo MRI (Echo Medical Systems, Houston, TX)

### Statistical Analysis

Sample sizes were determined based on our previous studies and preliminary results. All results were confirmed in at least two different batches of mice. Results from quantitative experiments were expressed as means ± SEM. GraphPad Prism 9.0 (San Diego, CA) was used for all statistical analyses. Where appropriate, significance was calculated by Student’s t test, one- or two-way ANOVA with Tukey’s or Sidak’s multiple comparison test.

## Acknowledgements

This work was supported by NIH/NIDDK grant DK114373, start-up funds from the University of Illinois at Urbana-Champaign, and Burnsides Laboratory Research Fund to B.W. The authors thank Karen Doty at the Comparative Biosciences Histology Laboratory and Lou Ann Miller at Materials Research Laboratory for technical support with histology analysis and electron microscopy analysis. Lipid Core at Vanderbilt University School of Medicine is supported by NIH grant DK020593.

---

**Fig. S1.**
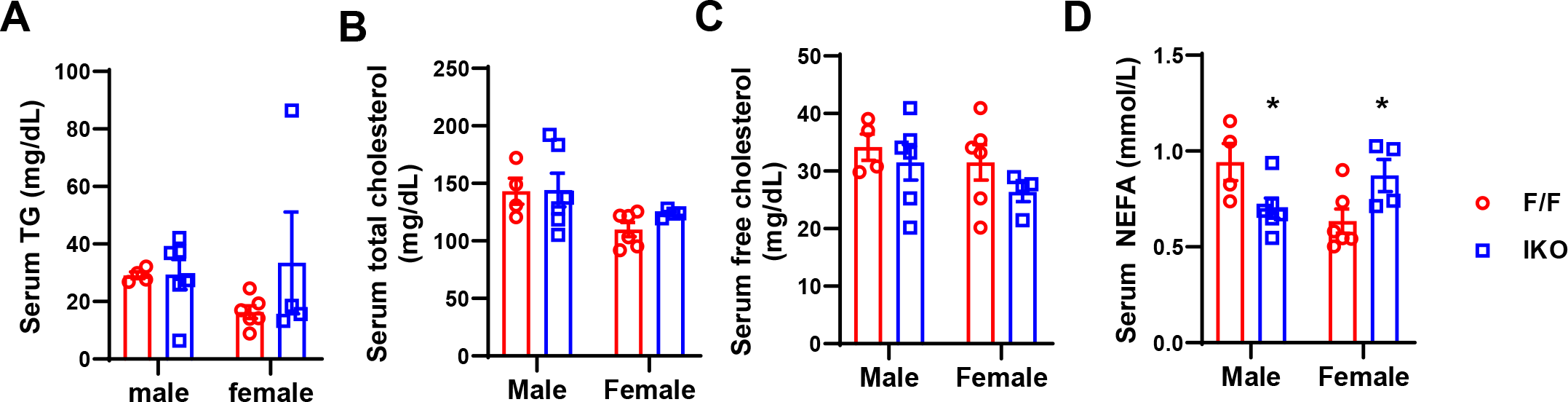
Serum lipids levels in chow diet fed Sec16b IKO and control mice. **A-D.** Serum TG (**A**), total cholesterol (**B**), free cholesterol (**C**) and NEFA (**D**) levels of chow diet fed Sec16b^F/F^ (F/F) and Sec16b^F/F^ Villin-Cre (IKO) mice (n=4-6/group). Mice were fasted 6 h before sacrificing. Values are means ± SEM. Statistical analysis was performed with Student’s t test. *P < 0.05.

**Fig. S2.**
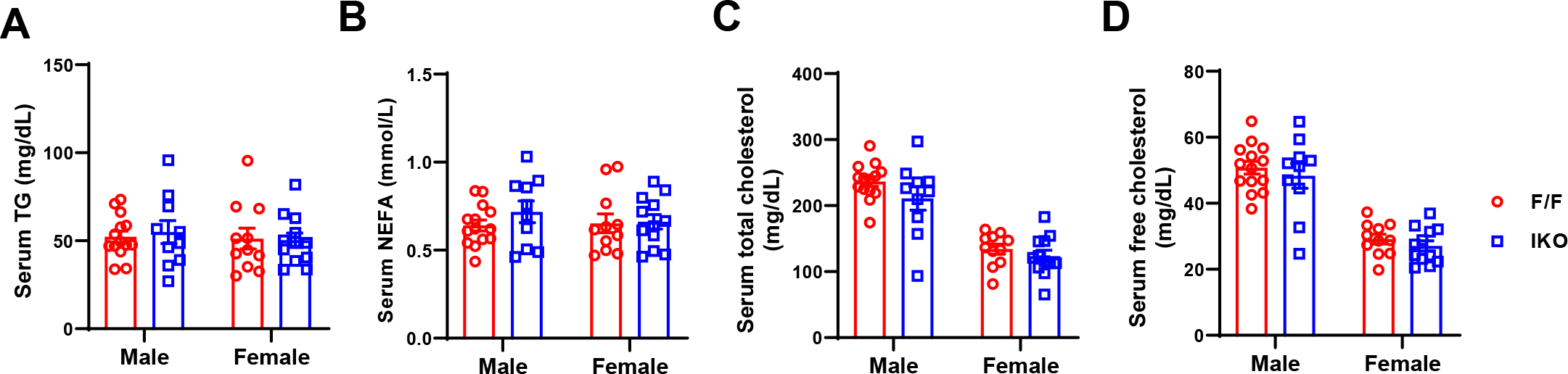
Serum lipid levels in HFD fed Sec16b IKO and control mice. **A-D.** Serum TG (**A**), NEFA (**B**), total cholesterol (**C**), and free cholesterol (**D**) levels of HFD fed Sec16b^F/F^ (F/F) and Sec16b^F/F^ Villin-Cre (IKO) mice (n≥10/group). Mice were fasted 6 h before sacrificing. Values are means ±SEM. Statistical analysis was performed with Student’s t test.

**Fig. S3.**
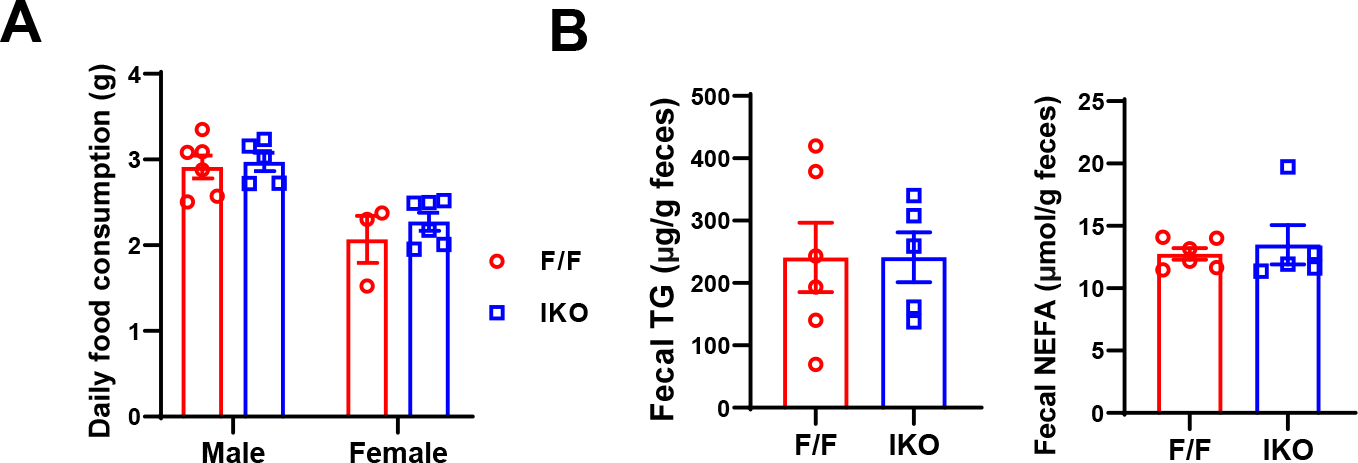
Food intake and fecal lipid levels in HFD fed Sec16b IKO and control mice. **A.** Daily food consumption of Sec16b^F/F^ (F/F) and Sec16b^F/F^ Villin-Cre (IKO) mice on HFD. Data was presented as average of four days (n=3-6/group). **B.** Fecal TG and NEFA levels of Sec16b^F/F^ (F/F) and Sec16b^F/F^ Villin-Cre (IKO) mice on HFD (n=5-6/group). Values are means ± SEM. Statistical analysis was performed with Student’s t test.

**Fig. S4.**
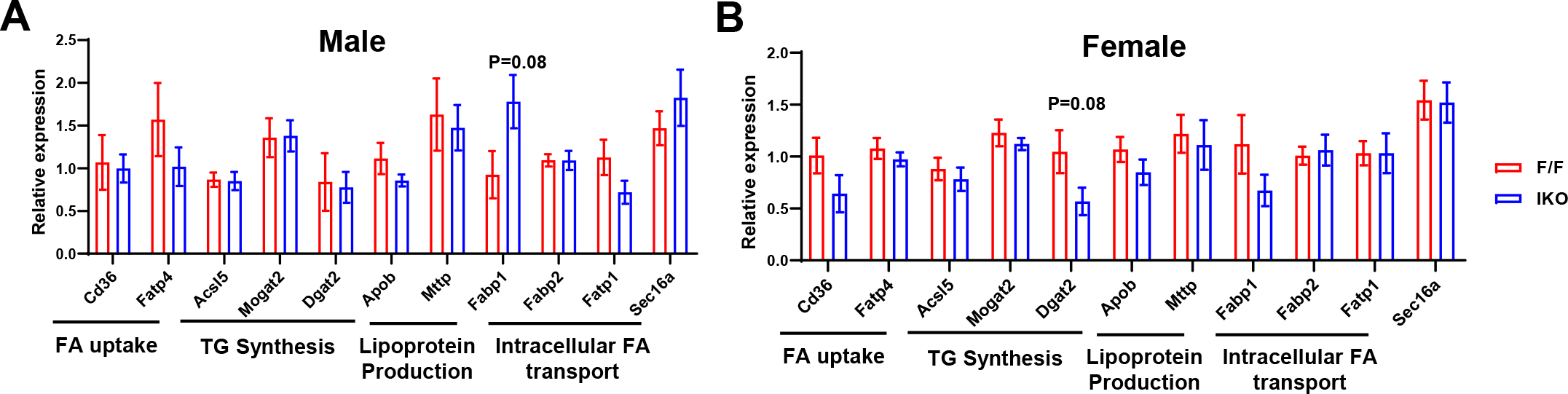
Gene expression in HFD fed Sec16b IKO and control mice. **A** and **B.** Gene expression in jejunum of male and female Sec16b^F/F^ (F/F) and Sec16b^F/F^ Villin-Cre (IKO) mice on HFD. (n= 4-6/group) Values are means ±SEM. Statistical analysis was performed with Student’s t test.

**Fig. S5.**
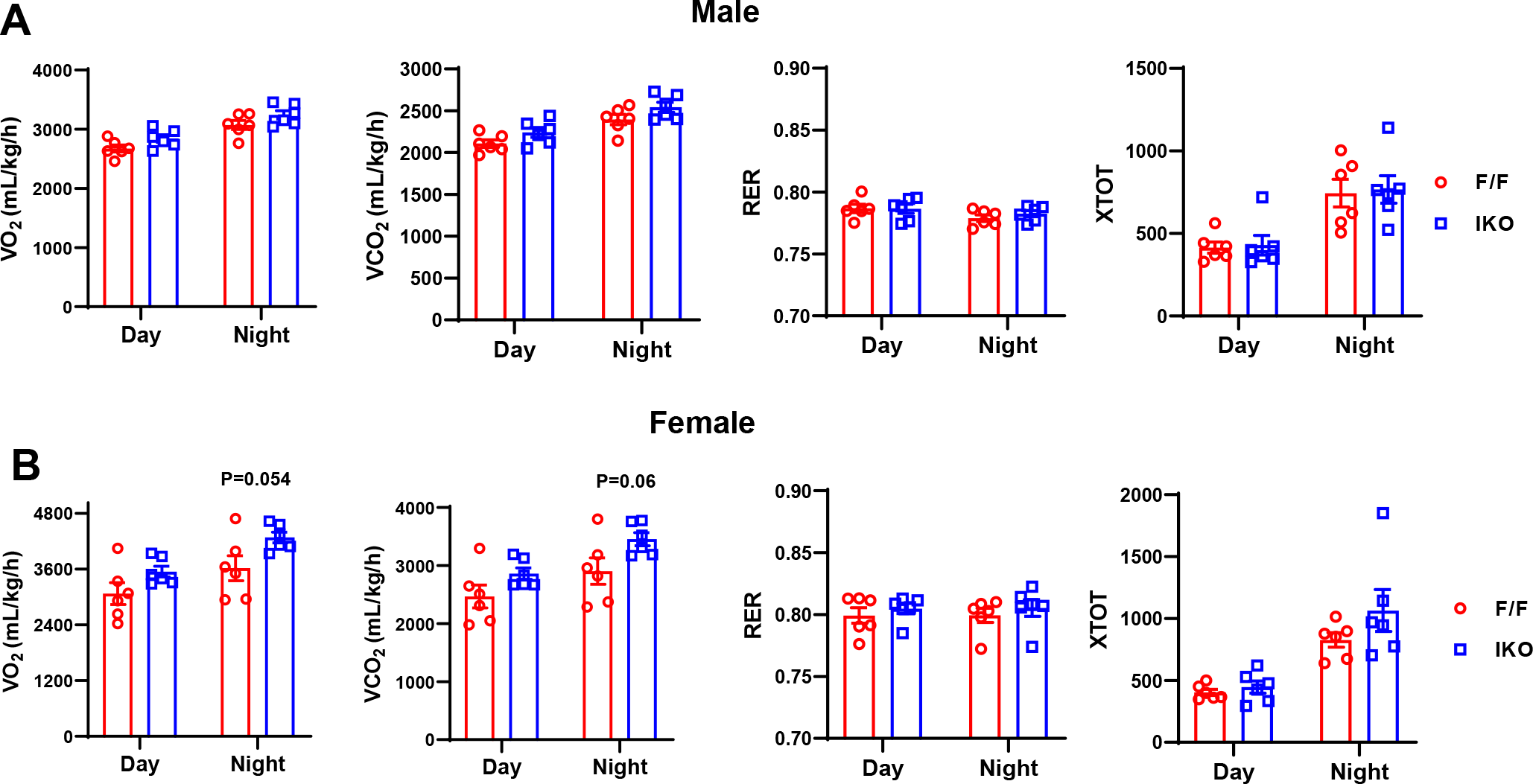
CLAMS analysis of HFD fed control and Sec16b IKO mice. **A** and **B**. CLAMS analysis of oxygen consumption rate, CO2 production rate, physical activity and respiration exchange ratio (RER) of Sec16b^F/F^ (F/F) and Sec16b^F/F^ Villin-Cre (IKO) miceon HFD (n=6/group). Values are means ± SEM. Statistical analysis was performed with two-way ANOVA.

**Fig. S6.**
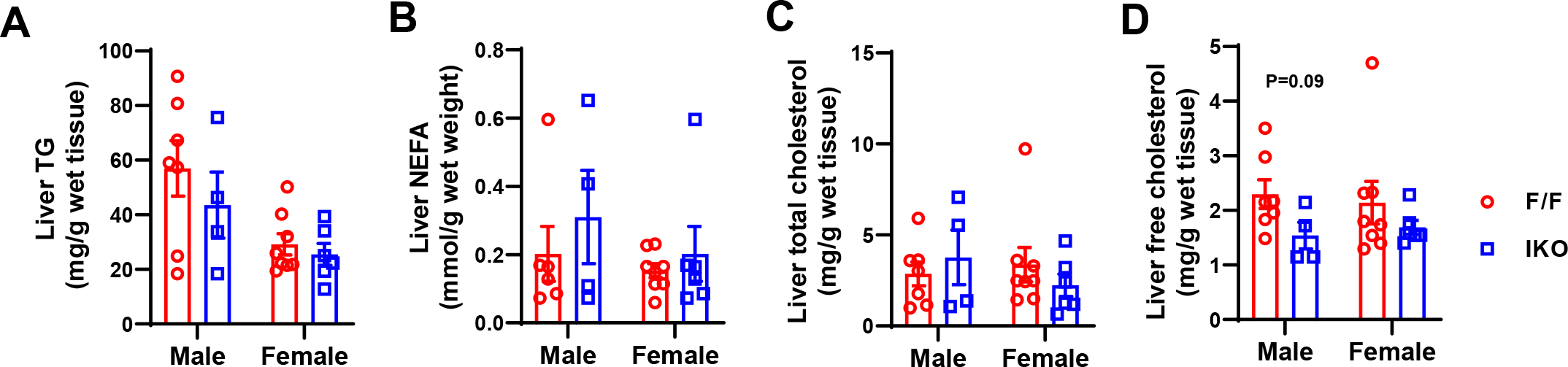
Hepatic lipid levels in HFD fed Sec16b IKO and control mice. **A-D.** Lipid levels in the livers of Sec16b^F/F^ (F/F) and Sec16b^F/F^ Villin-Cre (IKO) mice on HFD (n=4-7/group). Values are means ± SEM. Statistical analysis was performed with Student’s t test.

**Fig. S7.**
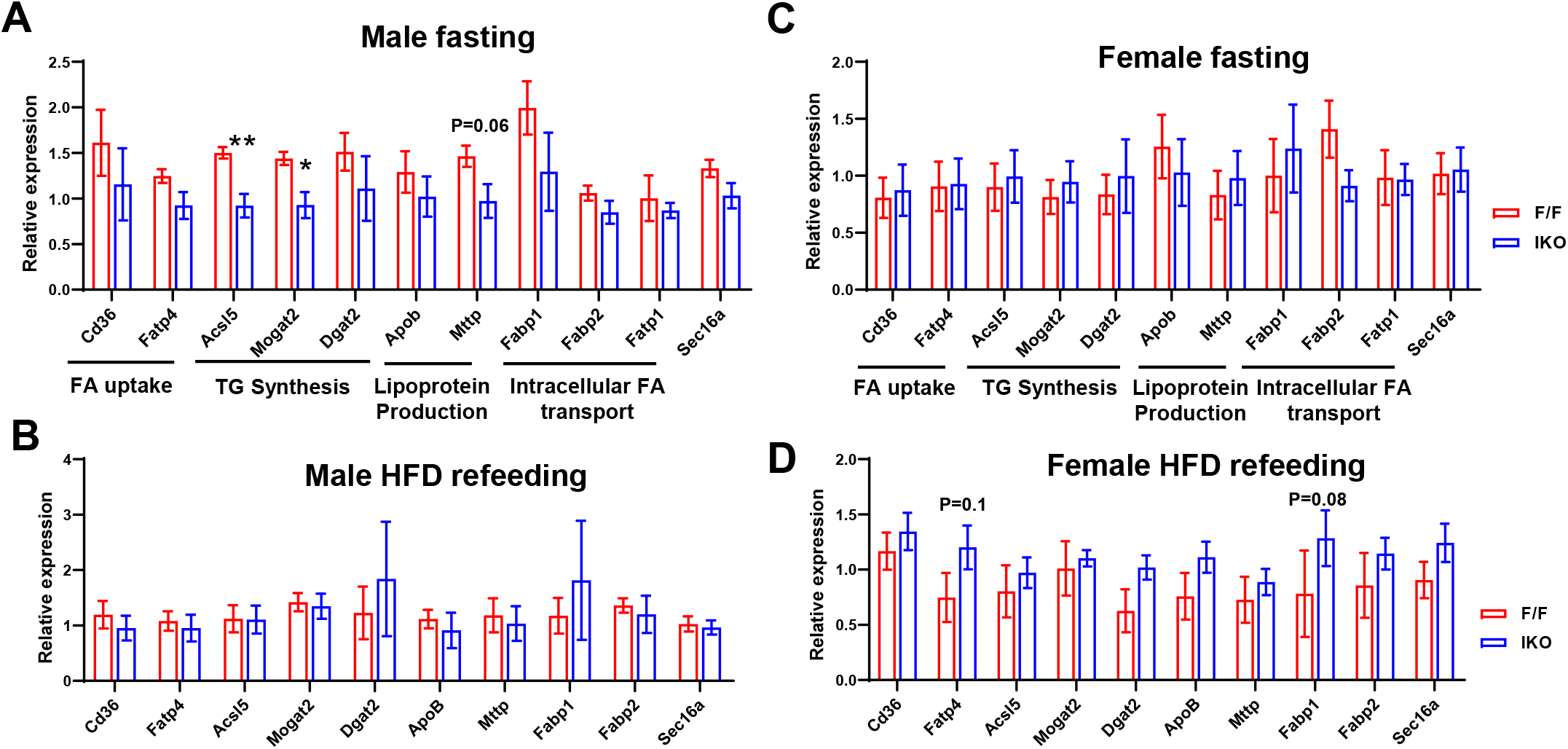
Gene expression in Sec16b IKO and control mice during fasting and HFD refeeding. **A** and **C.** Gene expression in jejunum of chow diet fed male (A) and female (C) Sec16b^F/F^ (F/F) and Sec16b^F/F^ Villin-Cre (IKO) mice after overnight fasting (n= 5-6/group). **B** and **D.** Gene expression in jejunum of male (B) and female (D) Sec16b^F/F^ (F/F) and Sec16b^F/F^ Villin-Cre (IKO) mice after HFD refeeding for 2 h (n= 3-5/group). Values are means ± SEM. Statistical analysis was performed with Student’s t test. *P < 0.05, **P < 0.01.

